## Supplementary information for "Label-free adaptive optics single-molecule localization microscopy for whole animals"

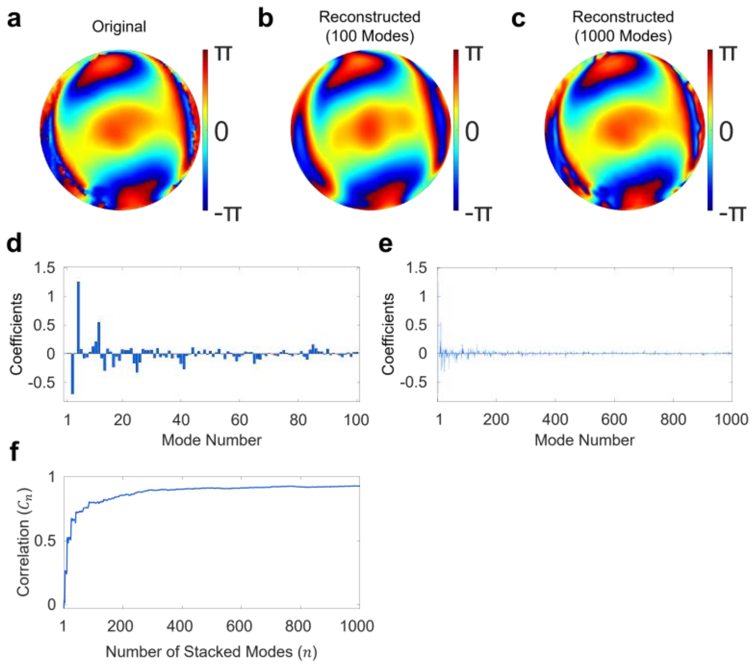

**Supplementary Figure 1. Aberration map decomposition by Zernike modes. a** Original aberration map identified by CLASS shown in the inset of Fig. 2c. **b** Reconstructed aberration map using the first 100 Zernike modes. It was quite similar to the original map in **a**, but slight differences were observed in the high spatial frequency region. Its correlation with the original aberration map was 0.797. **c** Same as **b**, but using the first 1000 Zernike modes. This aberration map was almost identical to the original one. Its correlation with the original aberration map was 0.926. **d**, **e** Coefficients of first 100 and 1000 Zernike modes constituting the reconstructed aberration maps in **b** and **c**, respectively. Note that the mode number over 20 still had non-negligible contributions. This indicates the complexity of the corrected aberration in this work considering that most of the previous reports dealt with aberrations composed of the first ~20 Zernike modes[1–4]. **f** Correlation $C_n$ between the original aberration map and the aberration maps reconstructed using first $n$ Zernike modes ($n = 1, 2, \cdots, 1000$). Correlation $C_n$ increases as the total number $n$ of Zernike modes used for reconstruction increases. Correlations $C_n$ for $n$ = 20, 100, 1000 were $C_{20}$ = 0.517, $C_{100}$ = 0.797, and $C_{1000}$ = 0.926, respectively.

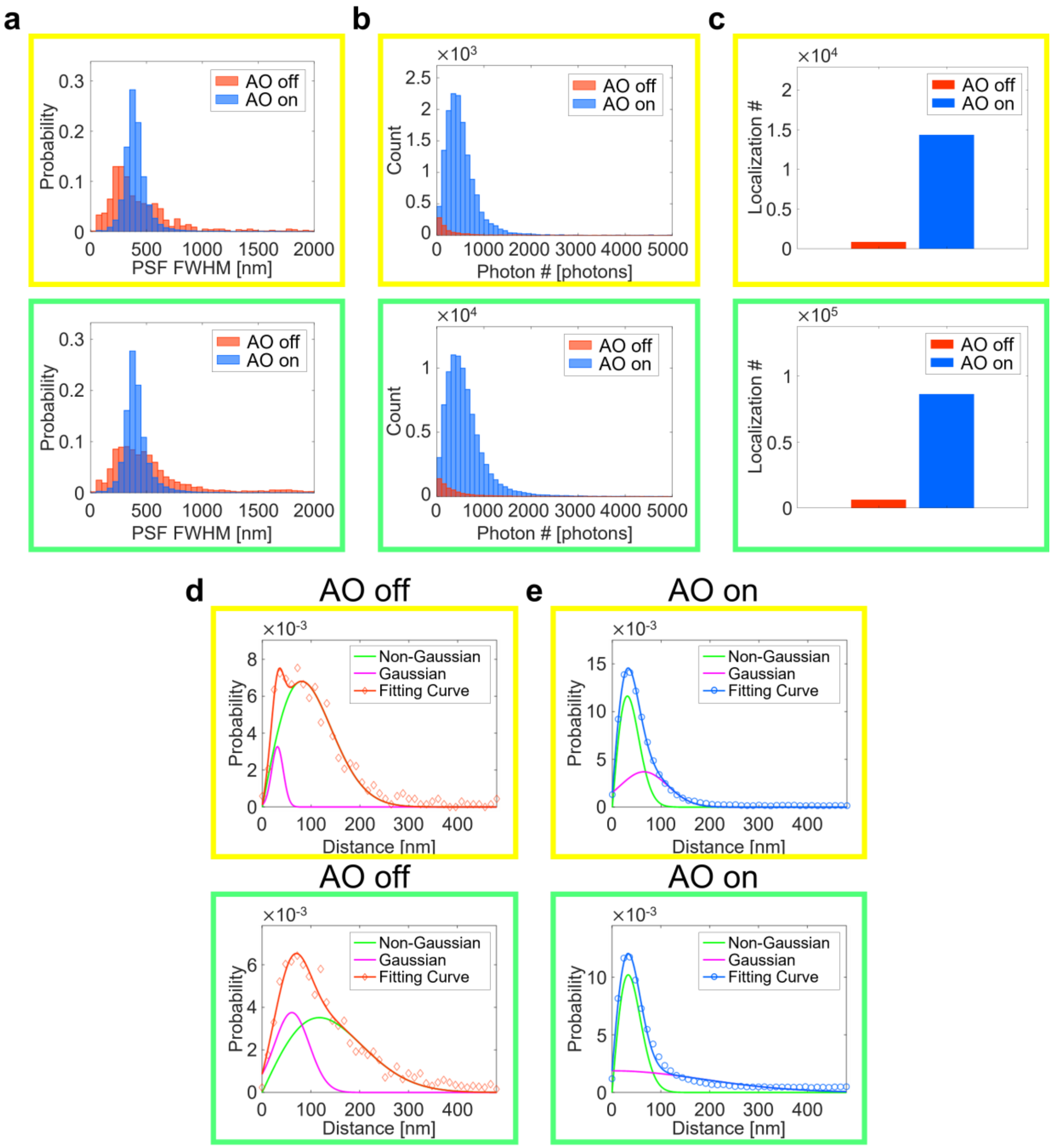

**Supplementary Figure 2. Comparison of SMLM quality metrics for the cell data in Figs. 2c, d.** Four different regions where single microtubules were isolated in the image field of view were selected as sparse regions (including the yellow box in Figs. 2c, d). Likewise, two different regions where microtubules were confluent were selected as dense regions (including the green box in Figs. 2c, d). Several parameters were extracted from these two types of regions for quantitative comparison. **a** Probability distribution of the FWHM of the PSFs identified by the ThunderSTORM. In both cases, it is noteworthy that PSF FWHMs were widely spread without AO, extended even beyond 1 μm. However, the distributions of PSF FWHMs were much narrower with AO. This supports that the PSF localization was robust with AO. **b** Histograms of photon number per emission PSF without and with AO. In the sparse region (yellow box), it increased from $5.68\times10^2$ photons to $1.08\times10^3$ photons (~1.90× increase). In the dense region (green box), it increased from $6.84\times10^2$ photons to $1.68\times10^3$ photons (~2.46× increase). **c** Bar graphs of the total number of localizations. There was a remarkable increase in the localization numbers with AO. In the sparse region (yellow box), it was increased from $7.88\times10^2$ to $1.43\times10^4$ (~18.1 × increase). In the dense region (green box), it was increased from $5.94\times10^3$ to $8.61\times10^4$ (~14.5× increase). **d**, **e** Probability density curves obtained by the nearest neighbor analysis of SMLM images without (**d**) and with (**e**)

AO as in Figs. 2j, l, but with separate fitting curves for a non-Gaussian and a Gaussian correction term (see Methods for the mathematical expression). With AO, the non-Gaussian curves, accounting for individual single molecules, dominantly contributed to the probability density curve in both sparse (yellow boxes in Figs. 2c, d) and dense (green boxes in Figs. 2c, d) regions (**e**). This indicates that AO worked well regardless of the density of cellular microtubules. Without AO, however, a different tendency was observed. In the dense region (green boxes in Figs. 2c, d), the Gaussian curve (magenta curve), describing the effect of different neighboring molecules, made a significant contribution. By comparing AO-off and -on raw images (Supplementary Video 1), it was visually confirmed that overlapping of PSFs among neighboring molecules was pronounced for AO off. This explains why most of microtubules were not recognizable in the dense region without AO (green box in Fig. 2g). In the sparse region (yellow boxes in Figs. 2c, d), the non-Gaussian curve was dominant as both cases of AO off and on (middle panel in **d**). However, the FWHM of the non-Gaussian curve, i.e., localization precision, was much broader without AO (57 nm) than with AO (22 nm).

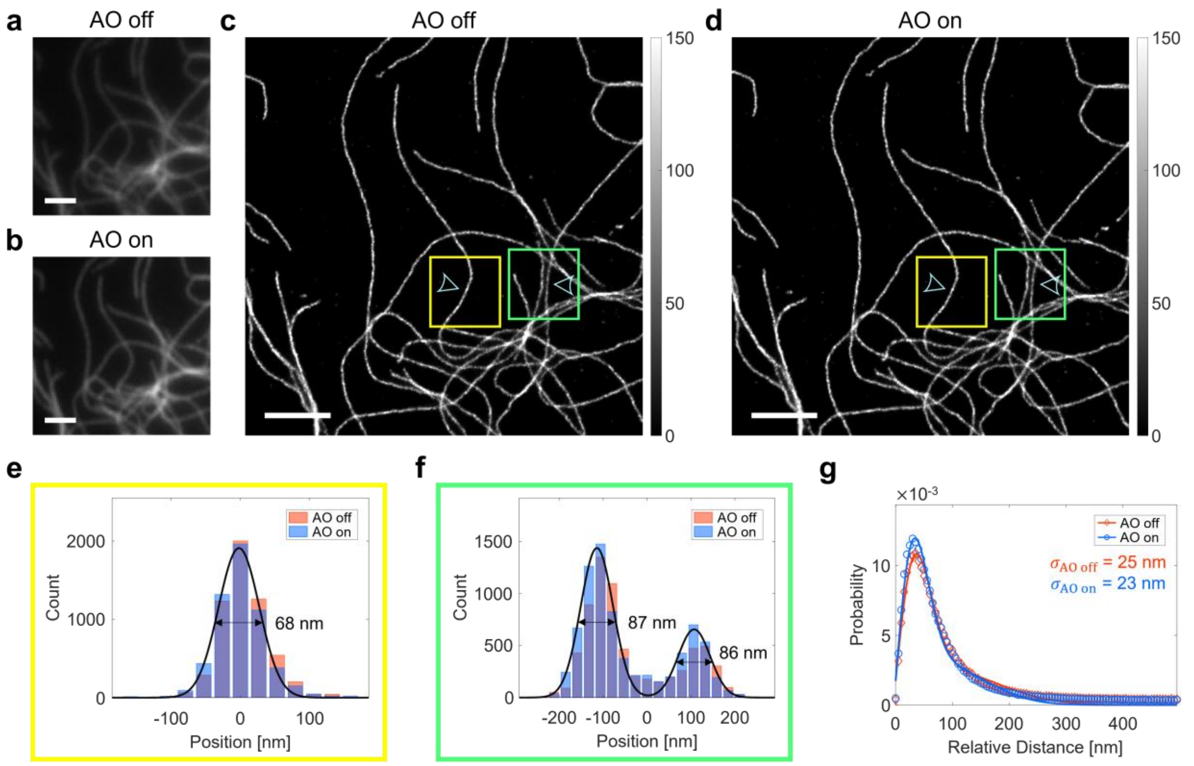

**Supplementary Figure 3. SMLM images of cells without an aberration layer. a**, **b** Diffraction-limited fluorescence images without (**a**) and with (**b**) AO, respectively. Images are normalized with respect to AO on. Scale bars indicate 2.5 μm. **c**, **d** SMLM images without (**c**) and with (**d**) AO, respectively. Here, only SMLM setup aberration was corrected when acquiring AO-on raw images of single-molecule blinking. Color bars indicate localization numbers. Scale bars indicate 2.5 μm. **e**, **f** Cross-sectional profiles at arrowheads in the yellow and green boxes in **c**, **d**, respectively.

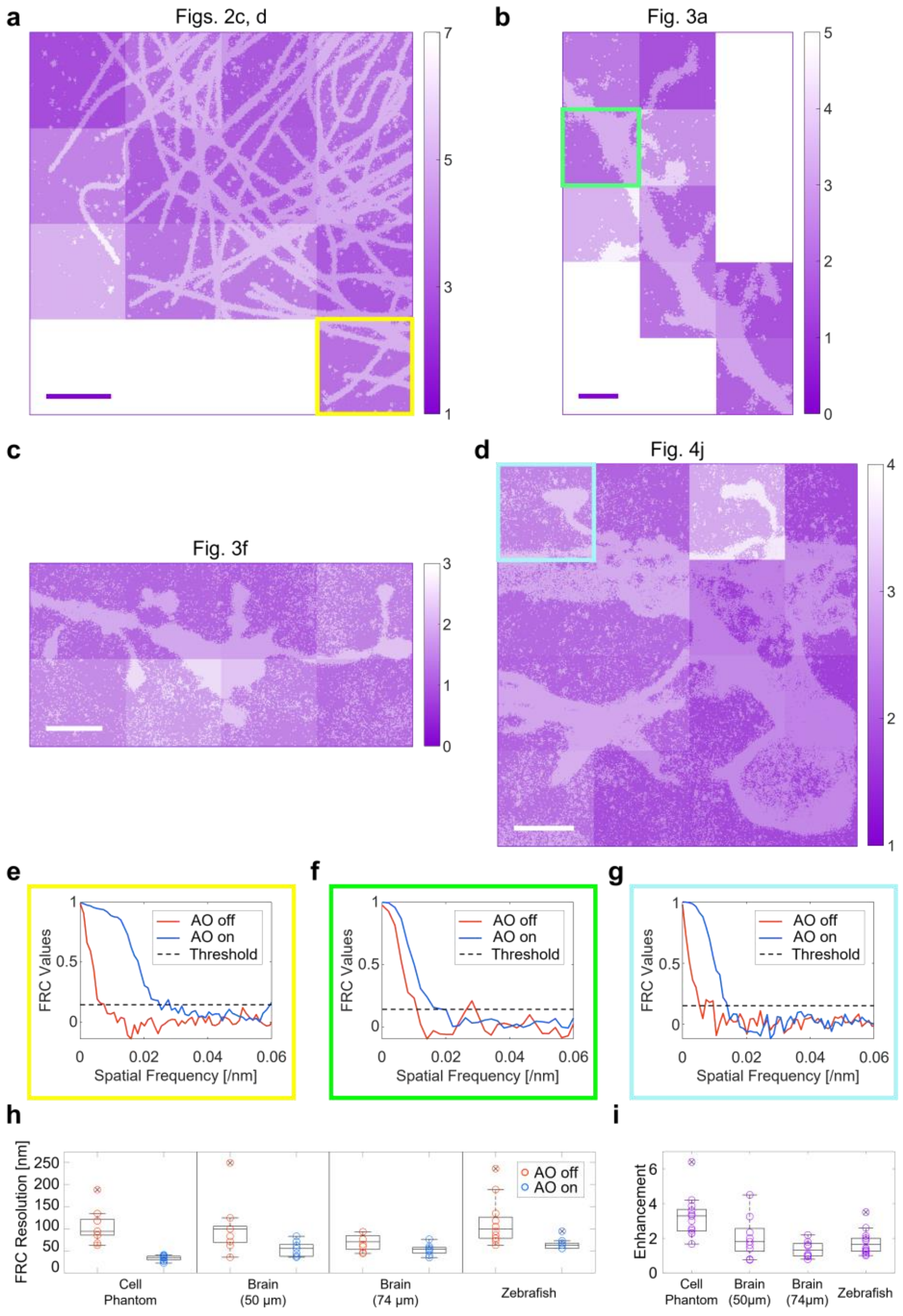

**Supplementary Figure 4. Fourier ring correlation (FRC) analysis of SMLM images. a-d** Enhancement of local FRC resolutions in Figs. 2c, 2d, 3a, 3f, and 4j, respectively. FRC resolutions were calculated as the reciprocals of cut-off spatial frequencies at which FRC value equals to 1/7[5]. Excluded areas are where no notable structures exist, which cause abnormally large values. Color bars indicate FRC resolution enhancements. Scale bars indicate 2.5 µm (**a**, **c**, **d**) and 2 µm (**b**). **e-g** Representative FRC spectra for the colored boxes indicated in **a**, **b**, and **d**, respectively. Nearest neighbor analysis curves in Figs. 3e, k showed mild distinctions between AO off and on though there were ~2-fold reductions in localization precisions with AO. However, FRC curve revealed an obvious broadening of FRC spectrum with AO. This is because FRC analysis incorporates the contribution of localization number as well as that of localization precision. **h** Box plots comparing FRC resolutions with and without AO. **i** Box plots of FRC resolution enhancements of the sub-areas in **a**-**d**. Outliers are marked with crosses. Note that AO-on FRC resolutions were well below 100 nm in all cases while many of AO-off resolutions were above 100 nm. This is consistent with the experimental observation that structures smaller than 100 nm were visible only with AO. Meanwhile, FRC resolution enhancements were largely similar in all local regions, suggesting that AO worked homogeneously everywhere.

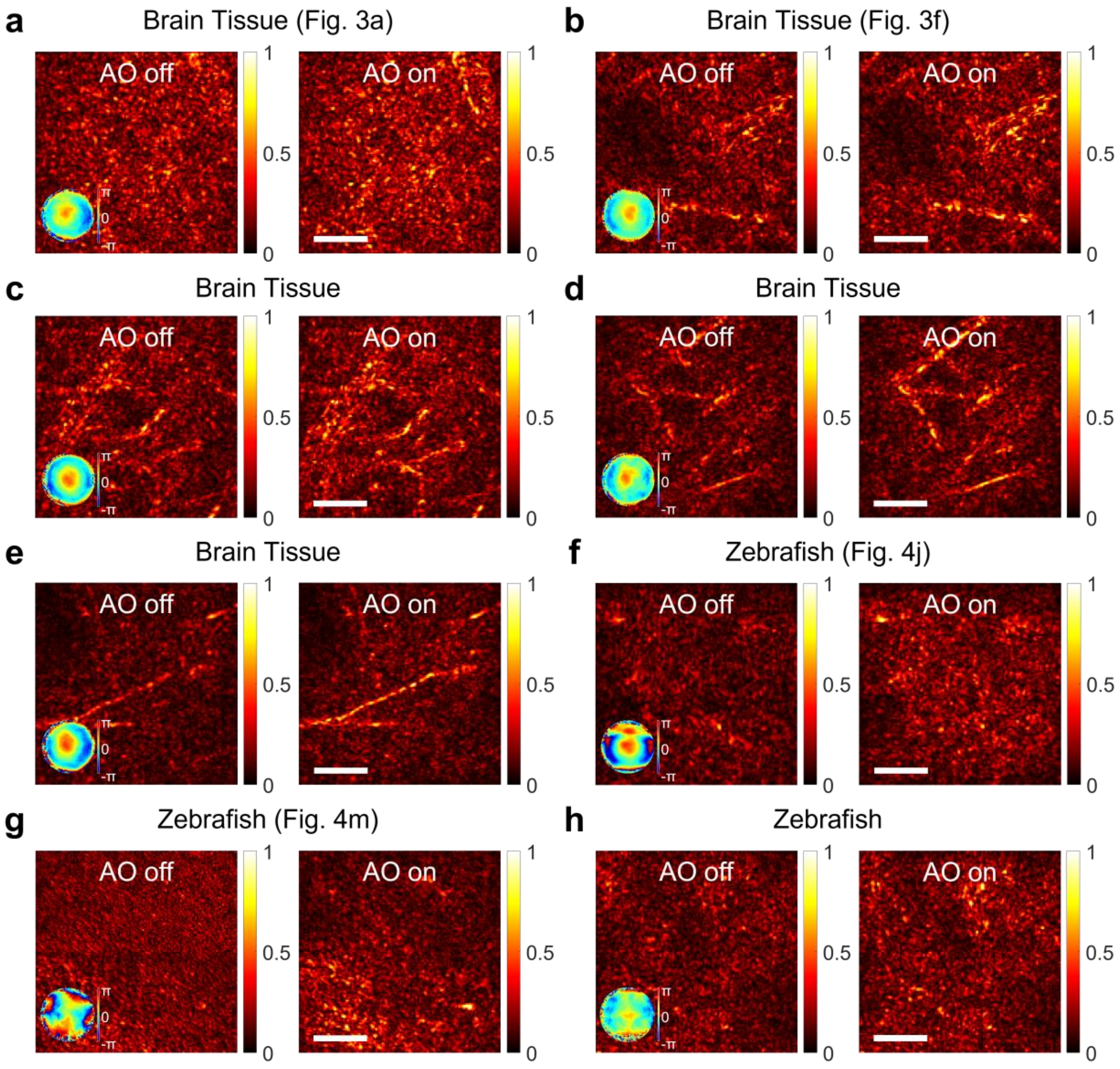

**Supplementary Figure 5. Reflection images of various samples. a-h** AO-off and -on reflectance images of various samples. Insets in each AO-off image indicate aberration correction maps. Color bars indicate normalized image intensity. Scale bars indicate 5 μm. In each sample, AO-off and -on images were normalized with AO-on image. Technically, AO-off images correspond to CASS images and AO-on images to aberration-corrected CASS images (Supplementary Note for detailed explanation on CASS and CLASS). Software-based AO via CLASS increased image intensity and SNR of CASS images. This effect of AO correction on the reflection images was much less dramatic than SMLM images (Figs. 3a, 3f, 4j, 4m). This is mainly because SMLM is much more prone to aberration than reflectance imaging due to its reliance on extremely weak single-molecule signals. In many cases, tissue aberrations were well resolved even when reflectance images hardly showed any recognizable structures. For example, some reflectance images looked like just speckle noises rather than specific biological structures as in **a**, **f**, **g**, and **h**. However, the same reflectance images were obtained for the repeated imaging of the same area. It suggests that these noise-like images originated from real sample structures. In some other cases, reflectance images revealed recognizable structures. For example, some of the dark, round patches observed in **b** and **e** might be blood vessels traversing in an out-of-plane direction. Blood vessels are quite frequently observed in LED transmission imaging. Sometimes, bright branch-like lines were observed (**b**-**e**). They are lipid-rich, highly reflective myelinated axons.

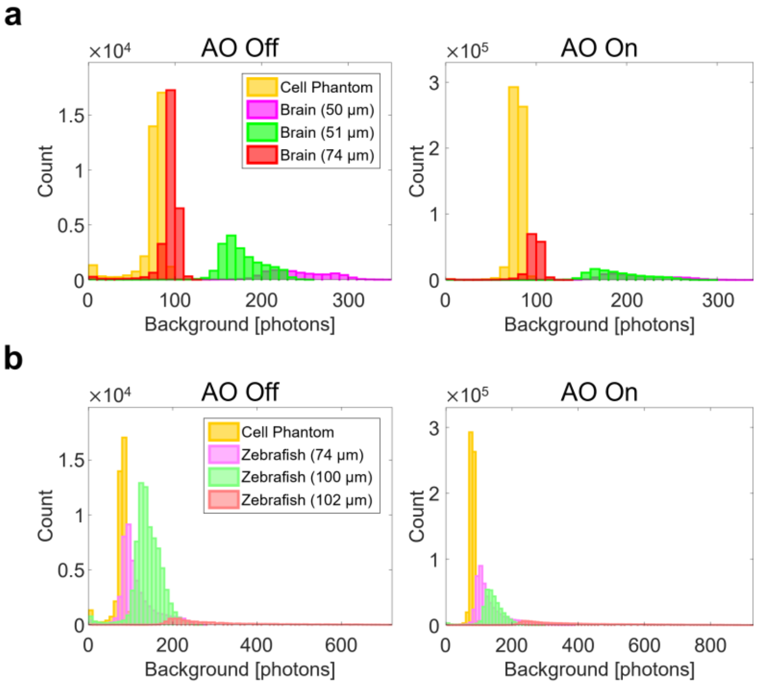

**Supplementary Figure 6**. **Histograms of background fluorescence for various samples.** **a**, **b** Background fluorescence per emission PSF obtained by ThunderSTORM analysis for a cell with an aberration layer, brain tissues (**a**), and intact zebrafish larvae (**b**). Cell phantom data was shown in both **a** and **b** for reference. Overall, background fluorescence was higher for brain tissues and intact zebrafish than cells. When imaging intact animals, the fluorescence background generally increases as the imaging depth increases (**b**). As a result, there is a tendency that localization precision and the localization number become lower at deeper depths.

| Sample | Figure | Depth [μm] | $A_{\mathrm{RMS}}$ | $w_{\mathrm{AO\ off}}$ [nm] | $w_{\mathrm{AO\ on}}$ [nm] | $\frac{S_{\mathrm{AO\ on}}}{S_{\mathrm{AO\ off}}}$ | $\sigma_{\mathrm{AO\ off}}$ [nm] | $\sigma_{\mathrm{AO\ on}}$ [nm] | $\frac{N^{\mathrm{loc}}_{\mathrm{AO\ on}}}{N^{\mathrm{loc}}_{\mathrm{AO\ off}}}$ |
|---|---|---|---|---|---|---|---|---|---|
| **Abrr.-Free Cell** | S3 | - | - | x: 551<br>y: 579 | x: 452<br>y: 494 | 1.01 | Whole Area | | |
| | | | | | | | 25 | 23 | 1.02 |
| **Cell + Abrr. Layer** | 2c, d | - | 2.22 | x': 1430<br>y': 1032 | x: 380<br>y: 444 | 4.27 | Whole Area | | |
| | | | | | | | 51 | 22 | 14.8 |
| | | | | | | | Yellow Box | | |
| | | | | | | | 57 | 22 | 25.5 |
| | | | | | | | Green Box | | |
| | | | | | | | 83 | 23 | 14.2 |
| **150-μm-Thick Brain Tissue** | 3a | 50 | 1.37 | x: 668<br>y: 668 | x: 413<br>y: 479 | 2.75 | Whole Area | | |
| | | | | | | | 66 | 38 | 9.32<br>Fig. 3b<br>9.80<br>Fig. 3c<br>6.75 |
| **200-μm-Thick Brain Tissue** | 3f | 74 | 0.983 | x: 744<br>y: 698 | x: 400<br>y: 475 | 3.92 | Whole Area | | |
| | | | | | | | 60 | 37 | 4.79<br>Fig. 3g<br>12.8<br>Fig. 3h<br>12.2<br>Fig. 3i<br>12.1 |
| **Zebrafish Cilia** | 4d | 82 | 2.14 | x': 1077<br>y': 1061 | x: 537<br>y: 517 | 1.93 | Whole Area | | |
| | | | | | | | 52 | 34 | 1.62 |
| **Zebrafish Commissural Tracts** | 4g | 100 | 3.08 | x: 783<br>y: 662 | x: 418<br>y: 489 | 1.17 | Whole Area | | |
| | | | | | | | 67 | 38 | 1.02 |
| **Zebrafish Nascent Myelin Sheath** | 4j | 52 | 2.13 | x: 619<br>y: 659 | x: 419<br>y: 434 | 3.34 | Whole Area | | |
| | | | | | | | 53 | 35 | 8.59<br>Arrowhead in Fig. 4j<br>37.4 |
| **Zebrafish Mauthner Axon** | 4m | 102 | 2.64 | x': 496<br>y': 2006 | x: 403<br>y: 458 | 8.86 | Whole Area | | |
| | | | | | | | Unmea. | 34 | 16.2 |

**Supplementary Table 1. Summary of measured parameters for various samples.** $A_{\mathrm{RMS}}$: RMS amplitude of sample aberration after removing trivial Zernike modes (tilt and defocus modes). $w_{\mathrm{AO\ off}}$ & $w_{\mathrm{AO\ on}}$: FWHMs of ensemble-averaged single-molecule PSFs for AO-off and -on, respectively. $S_{\mathrm{AO\ off}}$ & $S_{\mathrm{AO\ on}}$: Strehl ratios of ensemble-averaged single-molecule PSFs with AO-off and -on, respectively. $\sigma_{\mathrm{AO\ off}}$ & $\sigma_{\mathrm{AO\ on}}$: AO-off and -on localization precisions, respectively. $N^{\mathrm{loc}}_{\mathrm{AO\ off}}$ & $N^{\mathrm{loc}}_{\mathrm{AO\ on}}$: AO-off and -on localization numbers, respectively. Note that the x′- and y′-direction were defined as the longest axis and its orthogonal axis of the ensemble-averaged PSF, respectively. Also, $\sigma_{\mathrm{AO\ off}}$ was unmeasurable in Fig. 4m due to insufficient localization number.

### Supplementary Note

#### Details layout of experimental setup

This section describes the complete experimental setup, crucial alignment steps, and data acquisition procedures. The detailed layout of the experimental setup is shown in Supplementary Figure SN1. In the CLASS setup[6], we used a superluminal laser diode as a light source. Due to its short coherence length (~40 μm), it enables time-gated detection in the interferometry. Its wavelength was 678 nm, which lies near the center wavelength of fluorescence from Alexa Fluor 647 used for SMLM imaging. The 678-nm laser beam traveled through a 2-axis galvanometer mirror (GV). Note here that gray rectangles without name tags represent mirrors. The beam was then divided into two branches at BS1. One of them, called the sample beam (SB), was delivered to the sample through BS2, FM1, FM2, TL, and OL1 for widefield illumination. Then, SB retraced the same path back through OL1, TL, FM2, FM1, and BS2. Afterward, it traveled toward sCMOS camera through BS4. Meanwhile, the other beam, called the reference beam (RB), traveled toward the reference mirror through BS3. The reference mirror was attached to an automated translation stage for time-gated imaging by adjusting the reference beam path length. This was required because interference only occurs when the optical path length difference between SB and RB is within the coherence length. After being reflected from the reference mirror, RB retraced the same path back and traveled toward the grating through BS3. Passing through the grating, RB was diffracted into several branches. Among them, only the first-order branch was selected for oblique incidence of RB at the camera plane. The other branches were discarded by blocking with an iris. The first-order RB traveled toward the sCMOS camera through BS4. At the camera plane, SB and RB formed an interference pattern when the length difference between them was shorter than the coherence length.

The CLASS setup was designed for confocal reflectance imaging as well as interferometric reflectance imaging. Switching to confocal mode was accomplished by laying down the FM1, raising up BS5, and moving the sCMOS camera in front of OL2. The sCMOS camera was attached to a detachable magnetic base for freely changing its location. In confocal mode, RB splitted at BS1 traveled toward the sample through BS3, FM2, TL, and OL1 for point illumination. After being reflected from the sample, RB retraced the same path back and traveled toward the sCMOS camera through GV, BS5, and OL2.

In the SMLM setup, the activation and two excitation beams of wavelength 405 nm, 488 nm, and 647 nm, respectively, were coupled by BS6 and DM1. They traveled toward the sample through DM2, DM3, FM2, TL, and OL1 for widefield illumination. The emission beam from the sample traveled through OL1, TL, FM2, and DM3. It was then split into two branches at PBS1. One of them with a horizontal polarization was reflected from the SLM and phase-modulated for aberration correction. The other traveled toward the EMCCD through air without passing through the SLM. Both beams were combined by PBS2, traveling toward the EMCCD camera. They simultaneously arrived at spatially separated regions on the camera plane, enabling real-time comparison of AO-off and -on image frames.

The LED beam of a wavelength of 505 nm was used for low-magnification transmission imaging. It assisted in determining where to image. The back-illuminated LED beam passed through the sample and traveled toward a CCD camera through OL1, TL, FM2, DM3, and DM2.

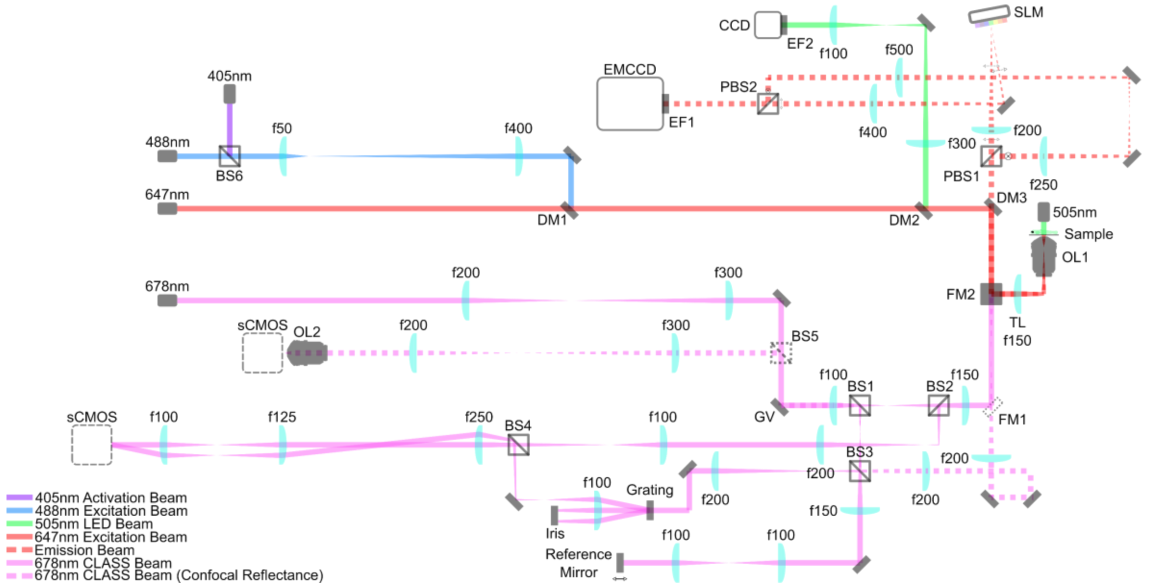

**Supplementary Figure SN1. Detailed layout of optical setup.** PBS1-2: polarizing beam splitters, BS1-6: beam splitters, FM1-2: flip mirrors, OL1-2: objective lenses, TL: tube lens, GV: 2-axis galvanometer mirror, DM1-3: dichroic mirrors, EF1-2: emission filters, gray rectangles without name tags: mirrors, and f$L_f$ ($L_f$ = 100, 125, …): lenses of focal $L_f$ in mm. One port of the commercial inverted microscope was devoted to CLASS imaging, and the other to SMLM imaging. Both setups share the sample, the objective (OL1), and the tube lens (TL). Also, they were switchable from each other using a built-in flip mirror (FM2) of the microscope. All lens pairs were used for either relaying images or adjusting beam size.

**Preparation of aberration correction**

As described above, the emission beam is split into two branches. For the fair comparison of AO-off and -on images, both images should have similar localization precision and photon number when there are no aberrations. To verify this, five different 200-nm-diameter fluorescent beads were imaged over 2,000 frames. Note that no aberrating layer was used in this test. For the test, the SLM was used as a mirror by setting all SLM pixels to the same phase values. Because SLM-unmodulated (“No SLM” in Supplementary Figure SN2) and -modulated (“Through SLM” in Supplementary Figure SN2) images were simultaneously recorded, ThunderSTORM analysis yielded 10 SMLM images for five beads. Half of them were constructed from SLM-unmodulated images, and the others from SLM-modulated images. Both SLM-unmodulated and SLM-modulated images yielded almost the same lateral localizations ($\sigma_x$ and $\sigma_y$) and photon numbers. This justifies a fair comparison of AO-off and -on images.

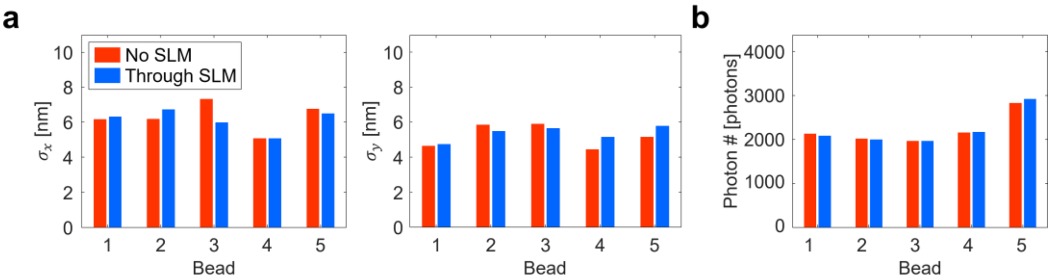

**Supplementary Figure SN2. Comparison of SLM-unmodulated and -modulated images when there were no aberrations. a**, **b** Bar graphs of lateral localization precisions (**a**) and photon number (**b**) for SMLM images of 5 different 200-nm-diameter fluorescent beads. 2000 frames were obtained for each bead. No SLM: SLM-unmodulated image, Through SLM: SLM-modulated image**.**

Next, we introduce a procedure to spatially match the aberration correction map displayed on the SLM with the objective pupil of the emission beam (Supplementary Figure SN3). The first step was to match their center positions. This process was accomplished by illuminating a mirror with a normally incident plane wave. Then, the excitation beam reflected from the mirror was focused on the center of the objective pupil. At the same time, the beam was focused on a certain position on the SLM display. That is to say, the focused beam position on the SLM is conjugated with the center of the objective pupil. To find this focused beam position, a grating pattern was displayed on the SLM (Supplementary Figure SN3a). On the grating pattern, we inserted a small square area with an equal phase value (Supplementary Figure SN3b). It served as a mirror patch. The pitch of the grating pattern was set fine enough that light impinging on it was diffracted away from the collection angle of the relay lenses, thereby unable to reach the camera view field. Therefore, the beam entering the mirror patch could only arrive at the camera, contributing to camera readout. The mirror patch was raster-scanned while recording the camera intensity (Supplementary Figure SN3c). The intensity was maximum when the mirror patch coincided with the focused beam indicated by a red circle in Supplementary Figure SN3b. That is, the center position of the aberration correction map was determined as the position of the mirror patch at which the intensity is maximum. In these measurements, the size of the mirror patch was set to ~100 μm. Too small or large mirror patch results in either negligible intensity variation or poor positioning precision.

The second step was to determine the diameter of the aberration correction map displayed on the SLM plane. To this end, confluent fluorescent beads were loaded on the sample plane. Then, the emission beam formed a solid circle on the SLM plane indicated by a red circle in Supplementary Figure SN3d. The diameter of the aberration correction map should be equal to that of this emission beam for precise correction. To measure its diameter, the scanning method explained above was used again, but a mirror slab was scanned instead of a mirror patch. In the beginning, a grating pattern was displayed on the SLM. Then, starting from the left side, the grating pattern was replaced by a mirror slab (Supplementary Figure SN3d). The mirror slab consisted of pixels with the same phase values. While expanding the mirror slab from left to right, the intensity of the camera image was recorded (Supplementary Figure SN3e). At some point, the emission beam started to overlap with the mirror slab. Then, the intensity began to increase. The intensity was saturated when the mirror slab fully covered the emission beam. The emission beam diameter could be evaluated as the length of the region where the intensity was increased.

Using the calibrated results explained above, the display region of the aberration correction map was determined (gray circle in Supplementary Figure SN3f).

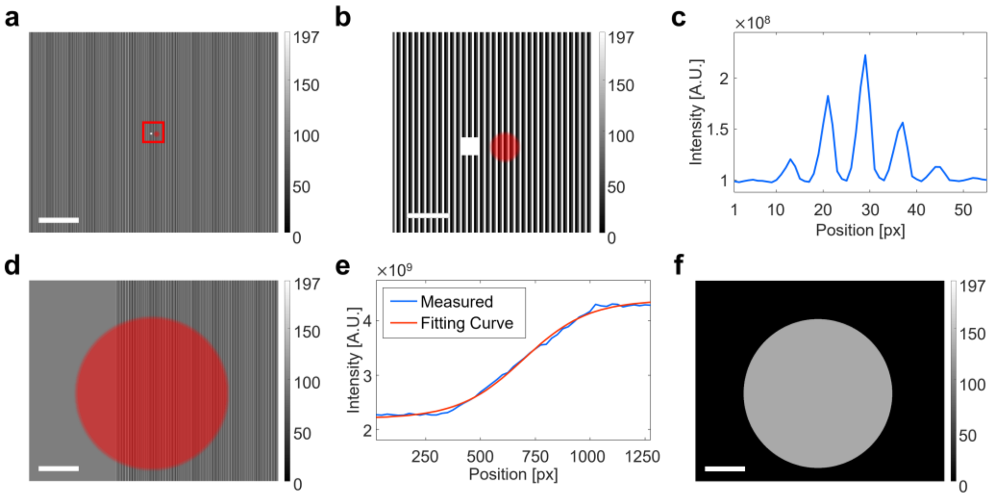

**Supplementary Figure SN3. Aberration correction map positioning procedure. a**, **b** Grating pattern with a small mirror patch displayed on the SLM (**a**) and a magnified view of red box (**b**). The white box is a small mirror patch. The solid red circle depicts the focused beam on the SLM plane. Color bars indicate SLM pixel values. Scale bars indicate 2.5 mm (**a**) and 250 μm (**b**). **c** Intensity of recorded image during mirror patch scanning. **d** Grating pattern with a mirror slab. The solid red circle is the emission beam on the SLM plane. Color bar indicates SLM pixel values. Scale bar indicates 2.5 mm. **e** Intensity of recorded image during mirror slab scanning. **f** Display region of the aberration correction map. A gray circle shows where correction maps to be displayed. Color bar indicates SLM pixel values. Scale bar indicates 2.5 mm.

The orientation and tilt of the aberration correction map displayed on the SLM were determined by a simple test (Supplementary Figure SN4). The correction map can be rotated counterclockwise by 0°, 90°, 180°, and 270° when displayed on the SLM. Also, it can be flipped horizontally. Hence, there are 8 cases in total. Note that vertical flip results in the duplicated cases because it is identical to 180° counterclockwise rotation and horizontal flip. Then, the correct combination can be found by trying to correct the aberration of fluorescent beads with an aberrating layer while displaying eight kinds of aberration correction maps. In our system, the aberration correction worked when the aberration correction map was counterclockwise rotated by 270° with no flip.

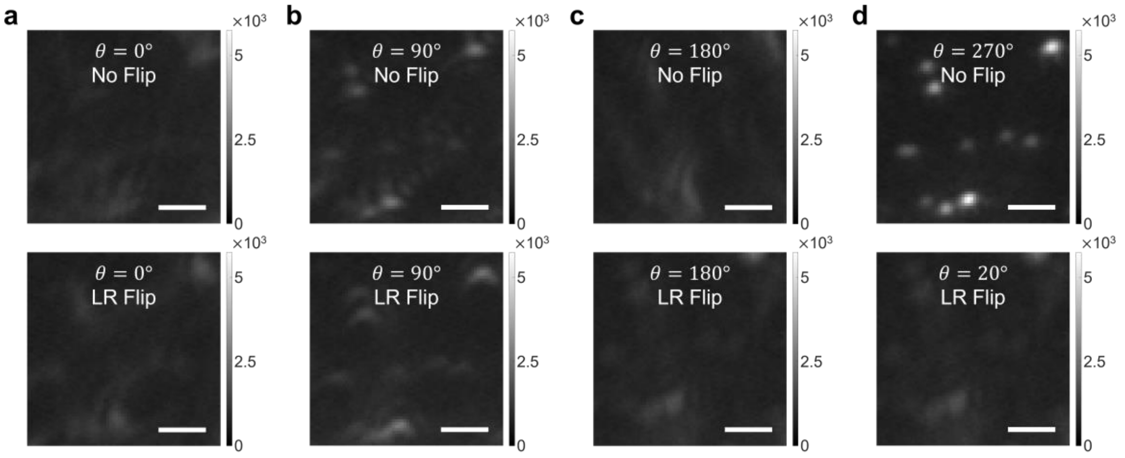

**Supplementary Figure SN4. Determination of orientation and tilt of displaying aberration correction map. a-d** Diffraction-limited fluorescence images of 200-nm-diameter beads for eight different orientation and tilt angles of the aberration correction map displayed on the SLM. Each was corrected by displaying the aberration correction map rotated by $\theta$ counterclockwise on the SLM ($\theta$ = 0°, 90°, 180°, and 270°). For each degree, the aberration correction map was displayed without (top row) or with a left-to-right flip (bottom row). Scale bars indicate 2 μm.

The optics layout of SMLM is different from that of the CLASS microscope except for the sample, the objective lens, and the tube lens (Supplementary Figure SN1). Therefore, it was necessary to separately correct for SMLM system aberration before correcting the sample-induced aberration identified by the CLASS (Supplementary Figure SN5). The system aberration $A$ of SMLM can be written as a linear combination of Zernike modes ($A = \sum_n c_n Z_n$). Here, $c_n$ is the coefficient of the $n^{\text{th}}$ Zernike mode $Z_n$. If each Zernike mode is displayed on the SLM while changing its coefficient, the aberration experienced by the light through the SLM changes according to the coefficient. Then, the PSF intensity is generally maximum when the aberration is minimum. Based on this property, the coefficient $c_n$ of each mode $Z_n$ consisting of the system aberration $A$ can be determined.

To this end, we tuned the coefficient $c_n$ of each mode $Z_n$ in such a way as to maximize the intensity of the PSF. Note that the first and second Zernike modes were excluded because they only spatially shift PSF rather than change its shape and intensity. Also, the PSF was indirectly measured by imaging a 200-nm diameter fluorescent bead. Starting from the Zernike mode 3, the measured PSF intensity was fitted with a double Gaussian function to find $c_3$ maximizing the PSF intensity (Supplementary Figure SN5a). While displaying $c_3 Z_3$ on the SLM, we added Zernike mode 4 and tuned its coefficient to find $c_4$ maximizing the PSF intensity (Supplementary Figure SN5b). The same process was repeated up to Zernike mode 20 to find SMLM system aberration (Supplementary Figure SN5c). Correcting system aberration increased PSF intensity and reduced PSF widths in both x- and y-direction (Supplementary Figure SN5d-e). With AO, PSF FWHM (Supplementary Figure SN5e) was slightly larger than the ideal FWHM $w_{\text{no abrr.}}$, which is calculated as

$$w_{\text{no abrr.}} \simeq 0.61 \frac{\lambda}{\text{NA}} = 0.61 \frac{670\ \text{nm}}{1.2} \simeq 341\ \text{nm}.$$

This suggests that weak SMLM system aberration remained even with AO. However, it was no trouble at all considering that sub-diffraction structures were well resolved with AO in our study.

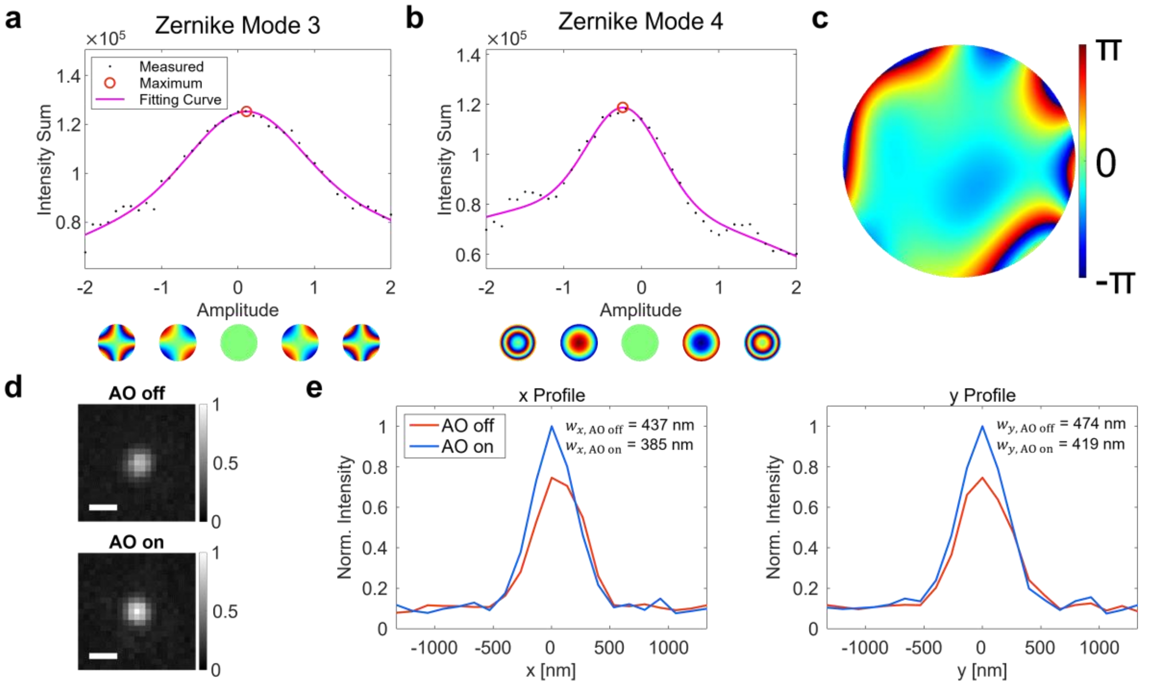

**Supplementary Figure SN5. SMLM system aberration correction.** **a**, **b** Intensity vs. mode coefficient plots for Zernike modes 3 (**a**) and 4 (**b**), respectively. **c** Identified SMLM system aberration. **d** Diffraction-limited fluorescence images of a 200-nm-diameter fluorescent bead without and with AO, respectively. Images are normalized with AO-on image. Color bars indicate normalized image intensity. Scale bars indicate 500 nm. **e** Line profiles of AO-off and -on images in **d**.

**Experimental procedure**

The imaging process of our AO-SMLM is described in Supplementary Figure SN6. For CLASS imaging, two sets of interference images were recorded (Supplementary Figure SN6a). They were composed of angle-scanned interference image sets of a mirror and the sample at the sample plane, respectively. Then, a reflection matrix $R$ was constructed using these two sets of images (Supplementary Figure SN6b). By applying the CLASS algorithm to $R$, we obtained a sample-induced aberration map (Supplementary Figure SN6c). The next step was checking the validity of the identified aberration map. It was accomplished by confirming that the software-based aberration correction well recovered the reflection image reconstructed from $R$ (Supplementary Figure SN6d). Then, the opposite phase of the calculated aberration was displayed on the SLM to correct the aberration experienced by the single-molecule emission PSFs.

After that, we acquired raw images of blinking single molecules (Supplementary Figure SN6e). As described above, the emission beam was split into two branches. One traveled through air without passing through the SLM and was recorded as AO-off raw images. The other traveled through the SLM and was recorded as AO-on raw images. Both AO-off and -on beams arrived at two different regions of the camera sensor for simultaneous recording. Lastly, both of these raw images were processed by ThunderSTORM software for constructing SMLM images and further post-processing (Supplementary Figure SN6f).

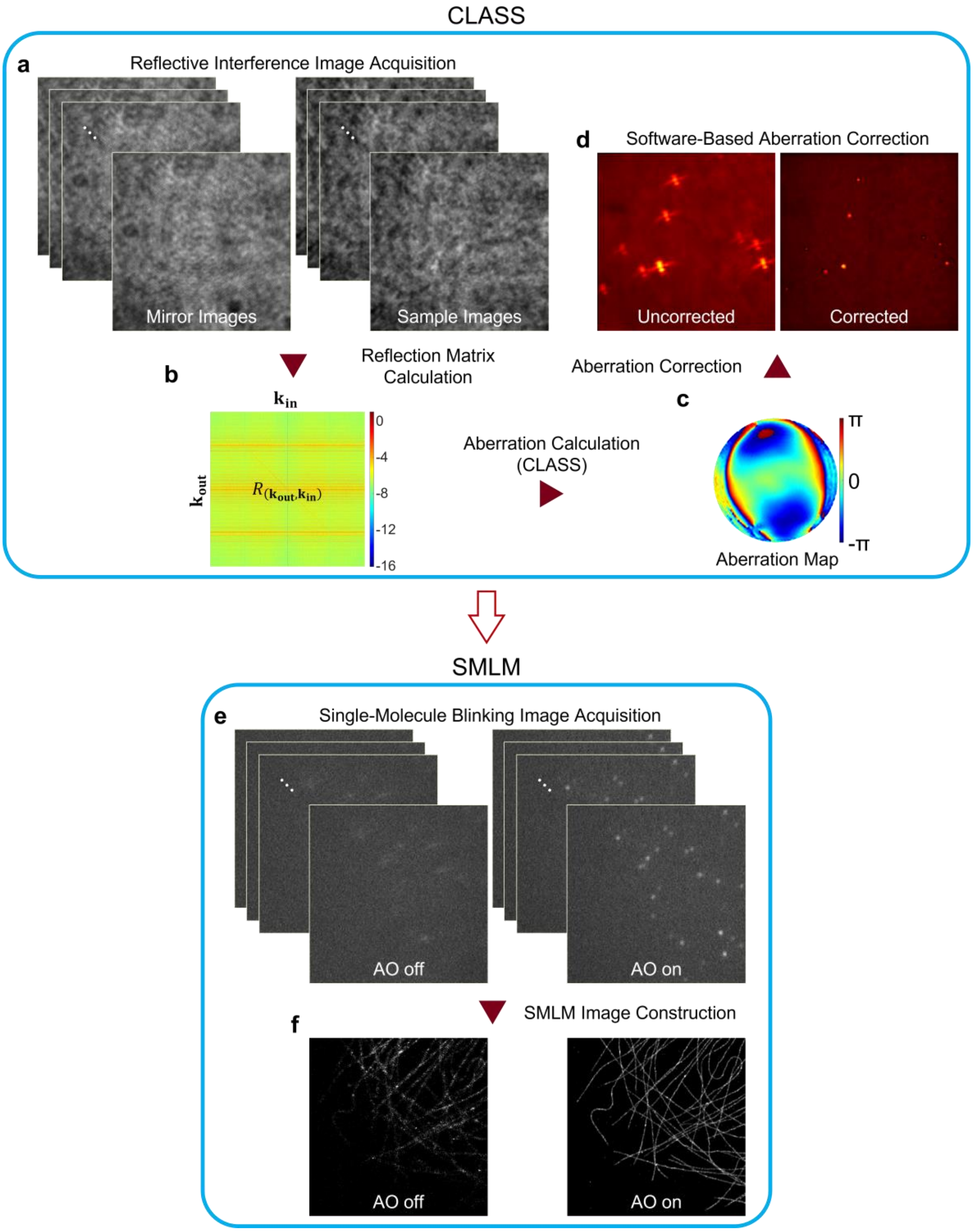

**Supplementary Figure SN6. Imaging process. a-d** CLASS microscopy process. **e**, **f** SMLM imaging process.

### Detailed process of CLASS algorithm

A rigorous description of CLASS starts from mathematically expressing interference. To this end, let $E_{\mathrm{S}}(\mathbf{r};\mathbf{k}_{\mathrm{in}})$ and $E_{\mathrm{R}}(\mathbf{r};\mathbf{k}_{\mathrm{in}})e^{-i\mathbf{k}_{\mathrm{G}}\cdot\mathbf{r}}$ be the electric fields of the sample beam (SB) and reference beam (RB), respectively, at position $\mathbf{r}=(x,y)$ on the camera plane. Here, $E_{\mathrm{S}}$ and $E_{\mathrm{R}}$ are complex numbers, and $\mathbf{k}_{\mathrm{in}}$ is the transverse wavevector set by illumination angle for each galvanometer mirror scanning position. The phase term $\mathbf{k}_{\mathrm{G}}\cdot\mathbf{r}$ is introduced because the RB is chosen from the first-order diffraction by a diffraction grating with spatial frequency $\mathbf{k}_{\mathrm{G}}$. Since RB is simultaneously rotated in sync with SB, the reference wave can be written as $E_{\mathrm{R}}(\mathbf{r};\mathbf{k}_{\mathrm{in}})=e^{i\mathbf{k}_{\mathrm{in}}\cdot\mathbf{r}}$. On the camera plane, interference occurs if the optical path length difference between SB and RB is smaller than the coherence length of the light source. Since a camera detects light intensity rather than the field itself, a camera image composed of SB and RB (Supplementary Figure SN7a) is written by the absolute square of their field sum. That is,

$$\begin{aligned} I(\mathbf{r};\mathbf{k}_{\mathrm{in}}) &= \left|E_{\mathrm{S}}(\mathbf{r};\mathbf{k}_{\mathrm{in}})+E_{\mathrm{R}}(\mathbf{r};\mathbf{k}_{\mathrm{in}})e^{-i\mathbf{k}_{\mathrm{G}}\cdot\mathbf{r}}\right|^2 \\ &= |E_{\mathrm{S}}(\mathbf{r};\mathbf{k}_{\mathrm{in}})|^2+|E_{\mathrm{R}}(\mathbf{r};\mathbf{k}_{\mathrm{in}})|^2+E_{\mathrm{S}}(\mathbf{r};\mathbf{k}_{\mathrm{in}})E_{\mathrm{R}}^*(\mathbf{r};\mathbf{k}_{\mathrm{in}})e^{i\mathbf{k}_{\mathrm{G}}\cdot\mathbf{r}}+E_{\mathrm{S}}^*(\mathbf{r};\mathbf{k}_{\mathrm{in}})E_{R}(\mathbf{r};\mathbf{k}_{\mathrm{in}})e^{-i\mathbf{k}_{\mathrm{G}}\cdot\mathbf{r}}. \end{aligned} \tag{1}$$

To extract the interference term $E_{\mathrm{S}}(\mathbf{r};\mathbf{k}_{\mathrm{in}})E_{\mathrm{R}}^*(\mathbf{r};\mathbf{k}_{\mathrm{in}})$, we take 2-D Fourier transform of the interference image (Supplementary Figure SN7b). We then select and crop the spectral component $\mathcal{F}\left\{E_{\mathrm{S}}(\mathbf{r};\mathbf{k}_{\mathrm{in}})E_{\mathrm{R}}^*(\mathbf{r};\mathbf{k}_{\mathrm{in}})\mathrm{e}^{i\mathbf{k}_{\mathrm{G}}\cdot\mathbf{r}}\right\}$ indicated by a red circle whose center is set by $\mathbf{k}_{\mathrm{G}}$. Cropped component is sent to a new coordinate system where its origin is $\mathbf{k}_{\mathrm{G}}$ (Supplementary Figure SN7c). This means that $\mathbf{k}_{\mathrm{G}}$ vanishes in the new coordinates. In this new coordinate system, the selected spectral component is written as $\mathcal{F}\{E_{\mathrm{S}}(\mathbf{r};\mathbf{k}_{\mathrm{in}})E_{\mathrm{R}}^*(\mathbf{r};\mathbf{k}_{\mathrm{in}})\}$. Taking the inverse Fourier transform, we obtain the $E_{\mathrm{S}}(\mathbf{r};\mathbf{k}_{\mathrm{in}})E_{\mathrm{R}}^*(\mathbf{r};\mathbf{k}_{\mathrm{in}})$ (Supplementary Figure SN7d).

The next step is to obtain $E_{\mathrm{S}}(\mathbf{r};\mathbf{k}_{\mathrm{in}})$ from the measured field $E_{\mathrm{S}}(\mathbf{r};\mathbf{k}_{\mathrm{in}})E_{\mathrm{R}}^*(\mathbf{r};\mathbf{k}_{\mathrm{in}})=E_{\mathrm{S}}(\mathbf{r};\mathbf{k}_{\mathrm{in}})e^{-i\mathbf{k}_{\mathrm{in}}\cdot\mathbf{r}}$. Removing the factor $e^{-i\mathbf{k}_{\mathrm{in}}\cdot\mathbf{r}}$ requires precise knowledge of $\mathbf{k}_{\mathrm{in}}$. This process is accomplished by imaging a scattering medium with the same incidence angles $\mathbf{k}_{\mathrm{in}}$. The scattered light by the scattering medium fully covers the pupil in the Fourier space (Supplementary Figure SN7e). Since this pupil rotates as the incidence angle $\mathbf{k}_{\mathrm{in}}$ changes, $\mathbf{k}_{\mathrm{in}}$ can be obtained from the center of the pupil. Then, the factor $e^{-i\mathbf{k}_{\mathrm{in}}\cdot\mathbf{r}}$ is removed in each measured spectrum $E_{\mathrm{S}}(\mathbf{r};\mathbf{k}_{\mathrm{in}})E_{\mathrm{R}}^*(\mathbf{r};\mathbf{k}_{\mathrm{in}})$ in Supplementary Figure SN7c by multiplying $e^{-i\mathbf{k}_{\mathrm{in}}\cdot\mathbf{r}}$.

We measure two sets of $E_{\mathrm{S}}(\mathbf{r};\mathbf{k}_{\mathrm{in}})$ for various $\mathbf{k}_{\mathrm{in}}$, one with a mirror on the sample plane and the other by placing the sample of interest, respectively. After that, two matrices, $P$ and $T$, are constructed to calculate the aberration (Supplementary Figure SN7f). The Fourier-transformed mirror image for the n$^{\mathrm{th}}$ incidence angle is vectorized and becomes the n$^{\mathrm{th}}$ column of a matrix $P$. Similarly, the vectorized Fourier-transformed sample image for the n$^{\mathrm{th}}$ incidence angle composes the n$^{\mathrm{th}}$ column of matrix $T$. Then the reflection matrix $R_{(\mathbf{k}_{\mathrm{out}},\mathbf{k}_{\mathrm{in}})}$ is obtained from the relation,

$$R_{(\mathbf{k}_{\mathrm{out}},\mathbf{k}_{\mathrm{in}})}\equiv T\times P^{-1}, \tag{2}$$

where $P^{-1}$ denotes the pseudoinverse matrix of $P$. The expression $R_{(\mathbf{k}_{\mathrm{out}},\mathbf{k}_{\mathrm{in}})}$ means that the row of $R$ correspond to incidence angle $\mathbf{k}_{\mathrm{in}}$ and their columns correspond to output angle $\mathbf{k}_{\mathrm{out}}$ (Supplementary Figure SN7g). Simply speaking, the $j^{\mathrm{th}}$ column of $R_{(\mathbf{k}_{\mathrm{out}},\mathbf{k}_{\mathrm{in}})}$ is a vectorized Fourier-transformed sample image for the $j^{\mathrm{th}}$ illumination angle $\mathbf{k}_{\mathrm{in}}^{(j)}$. $R_{(\mathbf{k}_{\mathrm{out}},\mathbf{k}_{\mathrm{in}})}$ shows every possible Fourier-transformed sample image for each illumination angle $\mathbf{k}_{\mathrm{in}}^{(j)}$. Here, $j=1,2,\ldots,N$ with $N$ the number of incidence angles.

The final step of CLASS is to process $R_{(\mathbf{k}_{\mathrm{out}},\mathbf{k}_{\mathrm{in}})}$ for calculating sample-induced aberration. For this, each column of $R$ is shifted by $\mathbf{k}_{\mathrm{in}}$ and add them all together. This coherent sum increases both the single-scattering signal preserving the sample information and the multiple-scattering noise. However, as images are coherently added, the single-scattering signal increases faster than the multiple-scattering one by a factor of the number $N$ of incidence angles. Inverse Fourier transform of this coherent sum gives an image whose single-scattering signal is significantly intensified (Supplementary Figure SN7g). This is the process termed collective accumulation of single-scattered waves (CASS)[7]. However, the CASS image still looks blurred (Supplementary Figure SN7h). This is due to the angle-dependent phase retardation known as an aberration. It undermines the proper accumulation of the single-scattering signal. In this example, the sample was intentionally defocused to introduce the well-known defocus aberration.

To correct the aberration, each column of $R_{(\mathbf{k}_{\mathrm{out}},\mathbf{k}_{\mathrm{in}})}$ is multiplied by a phase value ranging from 0 to $2\pi$. Then, the total intensity of the CASS image is maximized by adjusting phase values multiplied. Repeating this process for all the columns gradually increases the total image intensity. A similar process is made to the rows of $R_{(\mathbf{k}_{\mathrm{out}},\mathbf{k}_{\mathrm{in}})}$ to correct aberrations in the reflection beam path. After a few iterations, the intensity enhancement is saturated

(Supplementary Figure SN7i). This process is termed the closed loop accumulation of single scattering (CLASS) algorithm[8]. At this moment, aberration map is obtained by reshaping phase array applied to the rows of $R_{(\mathbf{k}_{\text{out}},\mathbf{k}_{\text{in}})}$ into a square form (Supplementary Figure SN7j). Finally, correcting this aberration yields an aberration-corrected CASS image (Supplementary Figure SN7k).

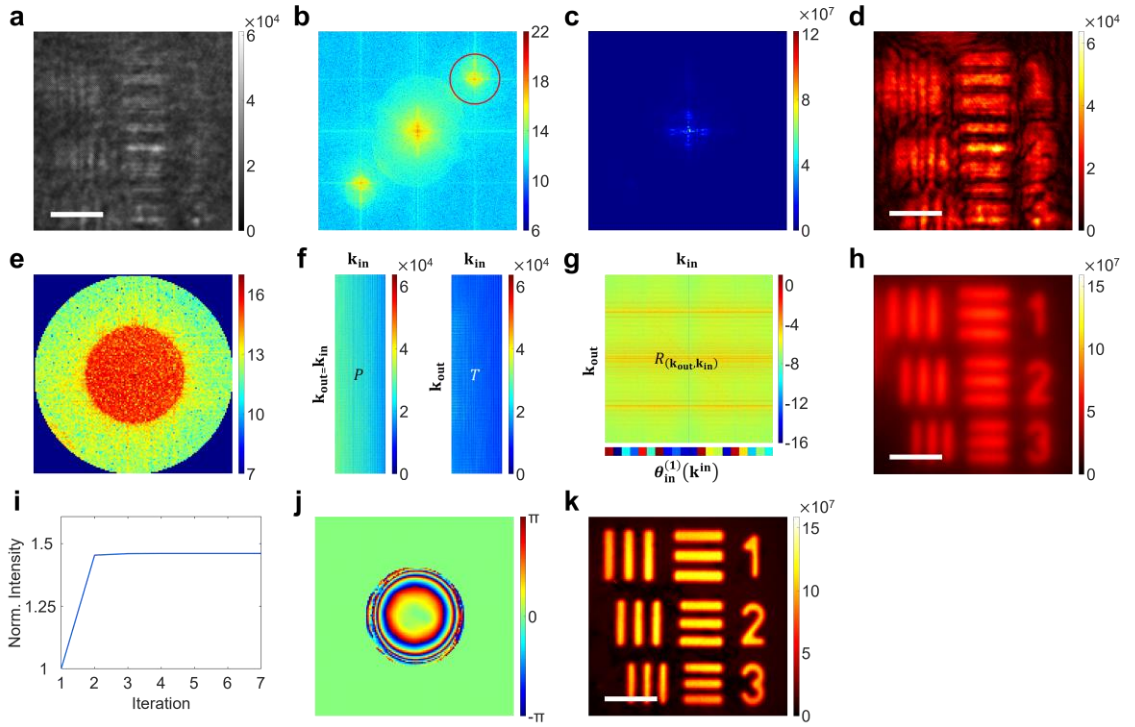

**Supplementary Figure SN7. Detailed process of CLASS microscopy. a** A raw interference image of a resolution target. Color bar indicates camera readout. Scale bar indicates 5 μm. **b**, **c** Fourier transform (**b**) of the interference image in **a** and selected spectral component (**c**) indicated by a red circle in **b**. Color bar indicates phase value. **d** Amplitude of the inverse Fourier transform of selected spectral component in **c**. Color bar indicates image intensity. Scale bar indicates 5 μm. **e** Pupil fully covered by multiple scattering induced by a scattering medium. Color bar indicates phase value. **f**, **g** *P* and *T* matrices (**f**) whose n$^{\text{th}}$ column is the vectorized Fourier-transformed image of a mirror and the sample, respectively. Reflection matrix *R* (**g**). Color bars indicates phase values. **h** CASS image. Color bar indicates image intensity. Scale bar indicates 5 μm. **i** Intensity enhancement during CLASS iteration. **j** Aberration map identified by CLASS. Color bar indicates phase value. **k** Aberration-corrected CASS image. Color bar indicates image intensity. Scale bars indicates 5 μm.
